# Duox activation in *Drosophila* Malpighian tubules stimulates intestinal epithelial renewal through a countercurrent flow

**DOI:** 10.1101/2023.10.18.562847

**Authors:** Zhonggeng Liu, Hongyu Zhang, Bruno Lemaitre, Xiaoxue Li

## Abstract

The gut must perform a dual role of protecting the host against toxins and pathogens while harboring mutualistic microbiota. Previous studies suggested that the NADPH oxidase Duox contributes to intestinal homeostasis in *Drosophila* by producing reactive oxygen species (ROS) in the gut that stimulate epithelial renewal. We find instead that ROS generated by Duox in the Malpighian tubules leads to the production of Upd3, which enters the gut and stimulates stem cell proliferation. We describe for the first time in *Drosophila* the existence of a countercurrent flow system, which pushes tubule-derived Upd3 to the anterior part of the gut and stimulates epithelial renewal at a distance. Thus, our paper clarifies the role of Duox in gut homeostasis and describes the existence of retrograde fluid flow in the gut, collectively revealing a fascinating example of inter-organ communication.

## Introduction

The digestive tract of animals forms a selective barrier that allows absorption of nutrients, ions, and water but limits contact with potentially damaging agents, such as toxins and pathogens. It also harbors a complex microbiota that contributes to host fitness through the breakdown of nutriments and the supply of vitamins (Thursby and Juge, 2017). The ability of the digestive tract to mount an effective immune response against pathogens while preserving the indigenous microbiota is enabled by specialized physical barriers and a sophisticated mucosal immune system (Sansonetti, 2004). In mammals, a plethora of innate and adaptive immunological mechanisms act in a regionalized manner along the digestive tract to ensure this selectivity. The efficiency of these mechanisms is supported by a strong ability of the gut epithelium to renew itself. Epithelium renewal through stem cell proliferation and differentiation repairs damages caused by pathogens, toxins, or by immunity itself, hence preserving the gut integrity (Allaire et al., 2018; van der Flier and Clevers, 2009). Rupture in this complex balance of immune and tolerance mechanisms in the digestive tract put the host at risk for infections, inflammatory diseases or gut leakage (Allaire et al., 2018; Buchon et al., 2013a; Sansonetti, 2004). The molecular mechanisms that ensure microbiota maintenance while preventing pathogenic infections remains largely unknown and remain difficult to tackle at the organismal level. Because of its anatomical and physiological similarity with the mammalian intestine, the *Drosophila* gut is a model of choice to study intestinal pathophysiology (Lemaitre and Miguel-Aliaga, 2013). Studies in *Drosophila* have already provided insights into mucosal innate immunity, intestinal sexual identity, epithelial renewal, host-commensal interactions, and more globally on how the gut is integrated at the organismal level (Colombani and Andersen, 2020).

Several mechanisms function in the *Drosophila* gut to restrict the growth of ingested pathogenic bacteria. Primary mechanisms include mechanical and chemical defenses. First, a chitinous semi-permeable barrier, the peritrophic matrix, is continuously produced in the anterior midgut and compartmentalizes the gut transversally into two spaces: the endoperitrophic and the ectoperitrophic space (Lehane, 1997). Similar to the mucus in mammalian guts, the peritrophic matrix confines the food bolus and bacteria to the endoperitrophic space, facilitating the digestive process while preventing direct interaction of the midgut epithelium with transiting bacteria (Hegedus et al., 2009; Kuraishi et al., 2011). In the middle midgut, secretions by V-ATPases create an acidic region that kills most ingested bacteria (Li et al., 2016; Overend et al., 2016). Lastly, peristalsis paced by visceral muscles surrounding the gut continuously pushes undigested food materials and microbes downstream, favoring their excretion (Benguettat et al., 2018; Du et al., 2016).

In addition to these chemical and physical barriers, two inducible immune mechanisms control both pathogenic and symbiotic microbes in the gut: secreted antimicrobial peptides (AMPs) and reactive oxygen species (ROS) (Lemaitre and Hoffmann, 2007). Ingested bacteria activate the expression of several antimicrobial peptide coding genes as well as other immune effector genes in specific domains along the digestive tract (Buchon et al., 2009a; Tzou et al., 2000). This response is initiated when peptidoglycan released from bacteria is sensed by pattern recognition receptors and activates the Imd signaling pathway (Bosco-Drayon et al., 2012; Neyen et al., 2012). The gut antibacterial response is kept in check by several negative regulators of the Imd pathway, notably by enzymatic PGRPs that scavenge peptidoglycan, preventing excessive and deleterious immune activation (Bosco-Drayon et al., 2012; Paredes et al., 2011). Two ROS producing enzymes, the NADPH oxidases Duox and Nox, are involved in intestinal microbe control in *Drosophila*. Nox produces superoxide in response to lactic acid derived from commensal *Lactobacillus* and has been shown to stimulate the basal level of stem cell proliferation (Iatsenko et al., 2018; Jones et al., 2013).

Duox contributes to the production of highly microbicidal ROS such as HOCl in the midgut (Ha et al., 2005a, 2009). In absence of Duox, pathogenic bacteria such as *Pectobacterium carotovorum* (*Ecc15*) proliferate in the gut (Ha et al., 2005a). Interestingly, Duox activity is selectively activated upon the sensing of uracil nucleotides that appear to be released by pathogenic but not commensal bacteria (Lee et al., 2013). This role of Duox in controlling gut microbes has also been observed in other insects (Xiao et al., 2017; Yao et al., 2016).

Furthermore, NADPH-produced ROS also increase defecation of food-borne pathogens through activation of TrpA1 ion channels in enteroendocrine cells (Du et al., 2016).

ROS are not only microbicidal but can induce tissue insults. As such, they cause collateral damage to the intestinal barrier that must be swiftly repaired for the insect to survive. The current model proposes that Duox-produced ROS induce epithelial damage, which causes the release of secreted ligands of the Upd family. Upds subsequently stimulate stem cell proliferation through the activation of the JAK/STAT and EGF-R pathways (Buchon et al., 2009b, 2010; Jiang et al., 2009; Kim and Lee, 2014). Consistent with this model, silencing *Duox* reduces the level of stem cell proliferation upon oral infection with enteric pathogens (Buchon et al., 2009b). Rapid recovery from a bacterial infection is possible only when the immune response is coordinated with epithelial renewal to repair the damage caused by both the pathogens and the immune response itself. While the regulation of Duox has been extensively studied, its exact contribution to gut immunity remains elusive. Intriguingly, *Duox* is weakly expressed in the midgut, and the gut phenotypes upon *Duox* knockdown appear only when a ubiquitously expressed driver is used (Buchon et al., 2009b; Leader et al., 2018). Thus, it is still largely unclear in which cell type(s) Duox is required and how Duox-derived ROS contribute, directly or indirectly, to eliminating pathogens and triggering epithelium renewal.

In this study, we further characterize the role of Duox in intestinal homeostasis. In opposition with the most commonly accepted model, we show that Duox is not directly required in the midgut, but rather in the Malpighian tubules (MpT), the insect equivalent of mammalian kidneys. Surprisingly, our study reveals the existence of a countercurrent flow in the gut that flushes signals produced by MpT, and more specifically the ROS-induced cytokine Upd3, from the posterior towards the anterior part of the digestive tract. Oral bacterial infections increase the intensity of the countercurrent flow, thus improving MpT-gut inter-organ signaling to promote the early phase of gut epithelium renewal. Collectively, our study not only revisits the function of Duox in the gut but also reveals an unexpected role of countercurrent flow in mucosal immunology.

## Results

### ROS-mediated intestinal stem cell proliferation does not require Duox in the gut

The role of Duox in intestinal homeostasis was established based on the observation that silencing *Duox* by RNAi reduces stem cell proliferation upon bacterial infection (Buchon et al., 2009b). However, these results were obtained by silencing *Duox* ubiquitously using the *da-Gal4* driver, and did not fully demonstrate that Duox is required in the gut (Buchon et al., 2009b). To further characterize the role of Duox in gut homeostasis, we monitored the level of epithelium renewal after oral infection with *P. carotovorum* (*Ecc15*) in wild-type flies and flies expressing a *Duox* RNAi. As expected, oral bacterial infection with *Ecc15* increases epithelium renewal as indicated by the higher number of mitotic stem cells labelled by phospho-histone 3 (PH3) staining. We first confirmed that silencing *Duox* ubiquitously with the *da-Gal4* driver reduced the level of mitotic stem cells as previously described. However, a kinetic analysis showed that Duox was required only in the early phase of epithelial renewal at 16 hours post infection (hpi), as previously described by Buchon et al. (Buchon et al., 2009b). The number of PH3 positive cells at 48 and 72 hpi were similar between wild-type flies and *Duox* RNAi flies (Figure 1A). To characterize more precisely where Duox was required, we knocked down *Duox* using *Gal4* drivers that are expressed in different intestinal cell types, notably progenitors (stem cells and enteroblasts, *esg-Gal4*), enteroblasts (*Su(H)GBE-Gal4*), enteroendocrine cells (*voila-Gal4*) and enterocytes (*Myo-Gal4*), and monitored stem cell proliferation at 16 hpi. Surprisingly, none of these *Gal4*-*Duox* RNAi combinations affected stem cell proliferation after infection with *Ecc15* (Figure 1B). In contrast, knocking down the other *Drosophila* NADPH oxidase Nox either in progenitors or enterocytes reduced stem cell proliferation, consistent with previous studies (Jones et al., 2013; Patel et al., 2019) (Figure 1C). Although our data confirm the involvement of Duox in the early phase of infection-induced stem cell proliferation, they also suggest that, unlike Nox, Duox is not required in the gut as initially proposed.

**Figure 1.**
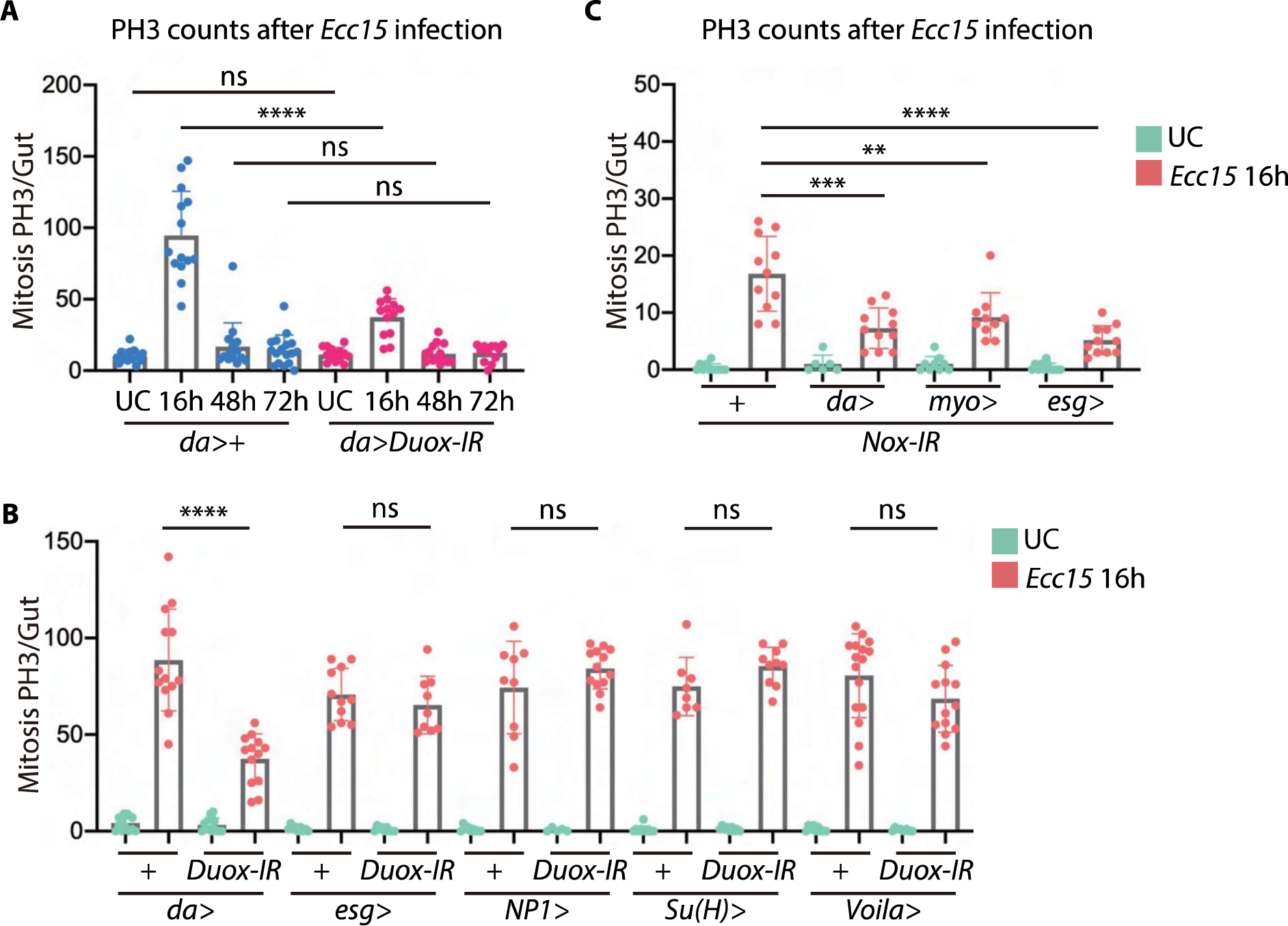
ROS-mediated intestinal stem cell proliferation does not require Duox in the gut (**A**) Counts of PH3-positive cells in the gut at different time points after infection show that ubiquitous (*da-Gal4*) RNAi silencing of *Duox* reduced mitosis at early time points after infection. (**B**) Counts of PH3-positive cells in the gut 16h after infection show that a reduced level of stem cell proliferation in the gut is observed only when silencing Duox with *da-Gal4* but not with driver expressing in intestinal progenitors (*esg-Gal4*), enteroblasts (*Su(H)GBE- Gal4*), enteroendocrine cells (*voila-Gal4*), or enterocytes (*Myo-Gal4*). (**C**) RNAi of *Nox* with ubiquitous (*da-Gal4*), enterocyte (*myo-Gal4*) and progenitor (*esg-Gal4*) drivers all reduce the level of epithelial renewal as shown by counts of PH3-positive cells. UC: unchallenged flies. ** for *P* between 0.001 and 0.01, *** for *P* between 0.0001 and 0.001, **** for *P*≤0.0001, ns: non-significant (A and B, Mann-Whitney test; C, one way ANOVA).

### Enteric infection induces a ROS burst in MpT through Duox

The MpT play an important role in osmo-regulation and waste removal, analogous to the mammalian kidney (Chapman et al., 2013; Dow et al., 1994a). These long tubules are connected to the gut at the midgut-hindgut junction, where fluid from MpT lumen is excreted into the digestive tract (Chapman et al., 2013) (Figure 2A). Publicly available transcriptomic studies indicate a much stronger expression of *Duox* in MpT compared to the gut (Leader et al., 2018). We assessed ROS production in the gut and MpT after *Ecc15* oral infection. To undertake these analyses, we stained both the gut and MpT using Dihydroethidium (DHE), a dye specific for superoxide, one of the products of Duox and Nox NADPH oxidases. DHE staining revealed that ingestion of *Ecc15* induced a ROS burst at an early time point (1.5 hpi) in MpT, but not in the gut (Figure 2B). Furthermore, the ROS level was much lower in the gut than in the tubules both before and after infection, suggesting tubules are the primary tissue of ROS production. We also confirmed this finding by measuring the level of hydrogen peroxide using a fluorometric hydrogen peroxidase assay (Ha et al., 2005a). We found that MpT produced more hydrogen peroxide than the gut upon infection, further confirming MpT as the main ROS production site (Figure 2C). We then investigated which NADPH oxidase was responsible for the ROS burst in MpT. Knocking down *Duox* but not *Nox* using the MpT principal cell-specific driver *uro-Gal4* significantly decreased ROS level in the tubules after infection (Figure 2D and E). Collectively, these data demonstrate that oral infection with *Ecc15* induces a ROS burst through Duox in MpT but not in the gut.

**Figure 2.**
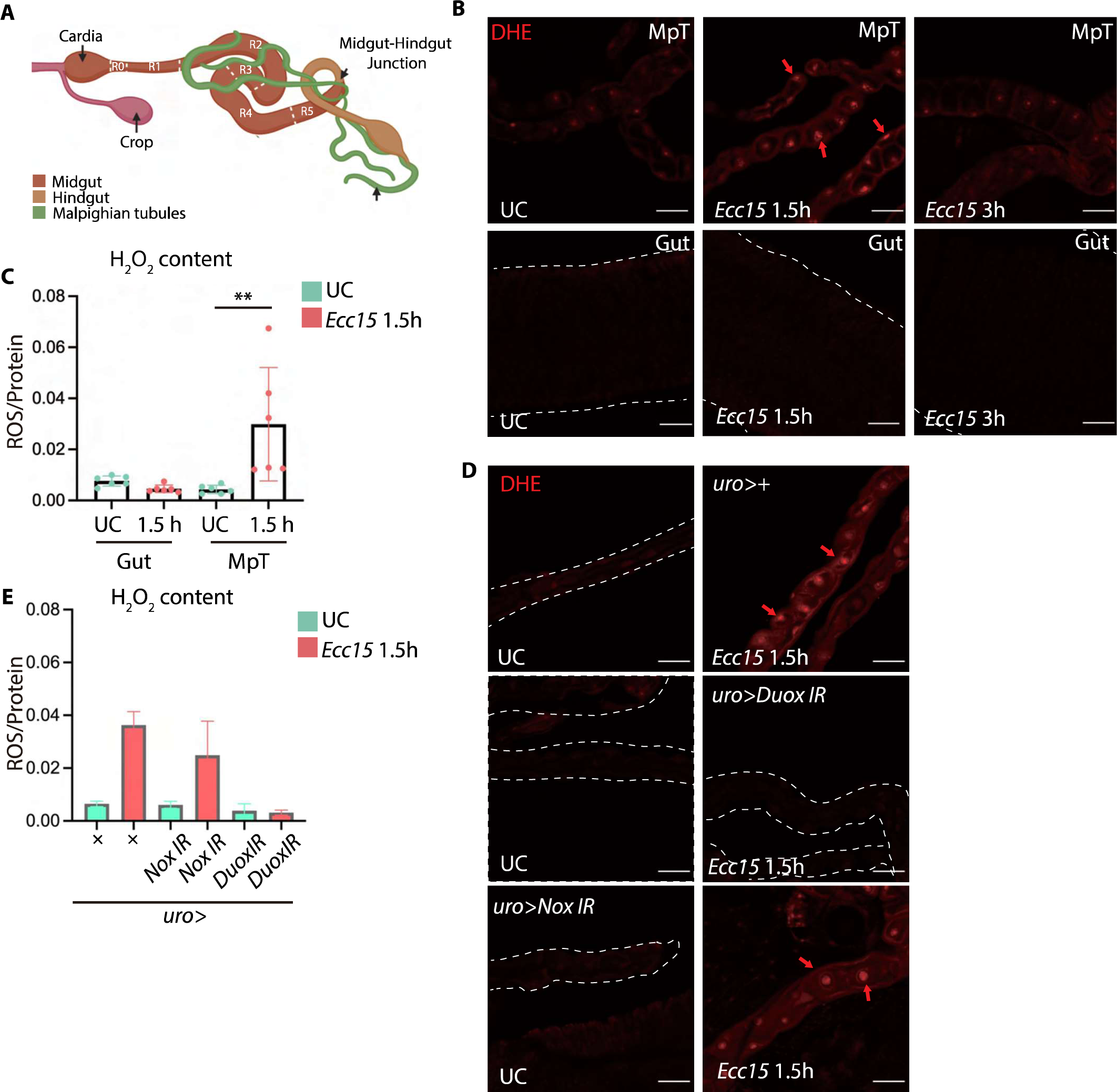
Enteric infection induces a ROS burst in MpT (**A**) A diagram of *Drosophila* digestive tract showing Malpighian tubules. (**B**) Confocal images of Malpighian tubules and gut stained with DHE. Malpighian tubules but not the gut show higher ROS level at 1.5h after oral infection with *Ecc15*. (**C**) Fluorometric measurement of hydrogen peroxide in Malpighian tubules of wild-type flies shows that a ROS burst occurs in the MpT but not the gut at 1.5h after *Ecc15* infection. (**D**) Confocal images of DHE staining of Malpighian tubules at 1.5hpi show that ROS is reduced by silencing *Duox* but not *Nox* in the MpT. (**E**) Fluorometric measurement of hydrogen peroxide in Malpighian tubules. UC: Unchallenged flies. ** for *P* between 0.001 and 0.01 with Mann-Whitney test, ns: non- significant.

### MpT derived Duox is necessary for gut homeostasis after infection

Our observations raise the intriguing hypothesis that stem cell proliferation in the midgut may depend on Duox function in the MpT rather than the gut. We therefore investigated whether ROS produced by Duox in the MpT contributed to stem cell proliferation after *Ecc15* infection. Strikingly, the number of mitotic intestinal stem cells was significantly reduced when *Duox* was silenced in MpT (Figure 3A) mimicking the results observed with the ubiquitous driver *da-Gal4* (Figure 1A). A kinetic analysis revealed that MpT Duox was required only in the early phase of epithelial renewal at 8 and 16 hpi. At the later time points (48 hpi and 96 hpi), there was no difference in the number of PH3 positive cells between wild-type flies and *Duox* knockdown flies (Figure 3A). Stem cell proliferation is associated with increased JAK-STAT pathway activity in the gut. Use of a *10XSTAT-GFP* reporter gene revealed that silencing *Duox* with the MpT *uro-GAL4* driver reduced JAK-STAT activity in the gut at 4 but not 16hpi upon *Ecc15* oral infection in the gut (Figure 3B).

**Figure 3.**
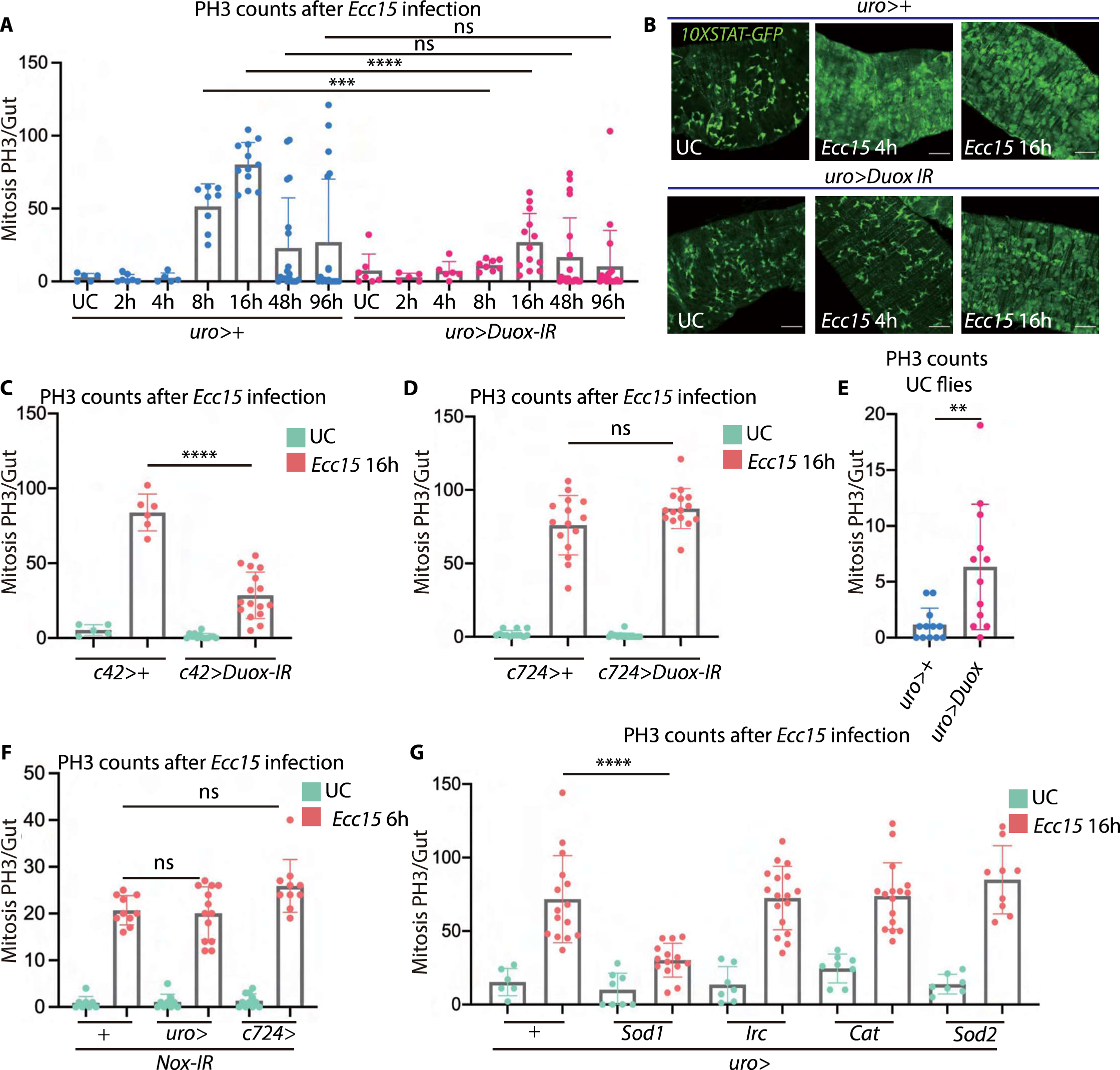
Duox is required in MpT to simulate intestinal epithelial renewal upon infection. (**A**) The level of epithelial renewal as monitored by PH3-positive cells count is reduced at early time points following *Ecc15* infection when *Duox* is knockdown in MpT (*uro-Gal4>Duox IR*). (**B**) Duox is required in Malpighian tubules for the early activation of JAK-STAT pathway activity in the gut after *Ecc15* oral infection as revealed using *10XSTAT-GFP*. (**C-D**) Duox is required in MpT principal cells but not stellate cells for gut epithelial renewal as revealed by specific knockdown with the principal cells *c42-Gal4* (C) and stellate cells *c724-Gal4* drivers (D). (**E**) *Duox* overexpression in Malpighian tubules (*uro-Gal4>uas-Duox*) promotes a low level of gut epithelial renewal in absence of infection. (**F**) *Nox* knockdown in either principal cells (*uro-Gal4>uas-Nox IR*) or stellate cells (*c724-Gal4>uas-Nox IR*) did not affect gut homeostasis after *Ecc15* oral infection. (**G**) Over expression of the cytosolic superoxide detoxifying enzyme gene *Sod1* in principal cells blocked *Ecc15* mediated gut epithelial renewal (*uro-Gal4>uas-Sod1*). Over expression of *Irc*, *Cat,* and *Sod2* did not impact gut epithelial renewal as monitored by PH3 count. UC: unchallenged flies. ** for *P* between 0.001 and 0.01, *** for *P* between 0.0001 and 0.001, **** for *P*≤0.0001, ns: non-significant (A, C, D and E, Mann-Whitney test; F and G, one way ANOVA).

The MpT epithelium consists of two types of cells with different roles in excretion: large principal cells and scattered stellate cells. Use of two additional *Gal4* drivers specific to the MpT principal cells (*c42-Gal4* and *capaR-Gal4*) and one distinct *Duox-IR* construct revealed that Duox is required in principal cells of MpT (Figure 3C and Figure S1A and B). Knocking down *Duox* in stellate cells, the other major MpT cell type, did not affect the level of epithelial renewal after *Ecc15* infection (Figure 3D). This indicates that Duox is required in principal but not stellate cells. As confirmation, removing Duox activity by tissue specific CRISPR knockout (Port et al., 2014) in the principal cells also blocked epithelial renewal (Figure S1C). Conversely, over-expression of *Duox* in MpT principal cells (*uro-GAL4*>*uas-Duox*) was sufficient to slightly increase the number of mitotic stem cells in absence of infection (Figure 3E). Silencing *Nox* in the MpT using either the *uro-GAL4* or the *c724-Gal4* did not affect stem cell proliferation upon oral bacterial infection (Figure 3F).

Antioxidant enzymes protect host tissues from the damaging action of ROS (Lambeth and Neish, 2014). In *Drosophila*, superoxide is detoxified by the cytosolic superoxide dismutase Sod1 and mitochondrial antioxidant enzyme Sod2, while hydrogen peroxide is detoxified by the extracellular catalase Irc and mitochondrial catalase Cat (Ha et al., 2005b; Lambeth and Neish, 2014). To further confirm that MpT ROS do contribute to intestinal stem cell proliferation, we over-expressed these four ROS scavenging enzymes in the MpT with the expectation that removing ROS in this organ would impact stem cell proliferation remotely in the intestine. We found that over-expressing *Sod1* but not the other three antioxidant enzymes in MpT reduced epithelium renewal upon oral infection (Figure 3G). Collectively, these data demonstrate that ROS produced by Duox in the principal cells of MpT control the early phase of gut epithelial renewal after oral bacterial infection.

### MpT derived Upd3 is necessary for gut homeostasis after infection

We then investigated the mechanism by which Duox in the MpT can remotely induce intestinal stem cell proliferation. A first question was to decipher whether Duox induces stem cell proliferation directly through ROS secretion or indirectly through a signaling molecule downstream of ROS. Stem cell proliferation is triggered upon activation of the JAK-STAT pathway by secreted proteins of the Unpaired family (Beebe et al., 2010; Buchon et al., 2009b; Jiang et al., 2009). In *Drosophila*, three unpaired proteins, Upd1, Upd2 and Upd3, contribute to JAK-STAT pathway activity by binding to the receptor Domeless (Agaisse et al., 2003; Osman et al., 2013). A recent study has shown that Duox activation in hemocytes leads to the production of Upd3 that remotely stimulates stem cell proliferation in the gut (Chakrabarti et al., 2016). This made Upd a good candidate signaling molecule to link Duox activity in MpT to intestinal epithelial renewal. We found that upon *Ecc15* infection, *upd3* but not *upd1* or *upd2* expression increases in MpT (Figure 4A). Interestingly, *upd3* gene induction was partially blocked upon *Duox* knockdown in the principal cells, indicating that its expression relies on Duox activity (Figure 4B). *upd3* induction was also blocked by over- expression of the cytosolic super oxide dismutase *Sod1* (Figure 4C). This indicates that, as observed in hemocytes, ROS produced by Duox is required for *upd3* induction in the MpT. We then investigated if *upd3* produced in MpT could remotely activate intestinal stem cell renewal. Strikingly, knocking down *upd3* in MpT principal cells using *uro-Gal4* reduced the number of PH3 positive cells in the gut upon infection (Figure 4D). Silencing *upd3* produced a reduction in stem cell proliferation comparable to silencing of *Duox* (Figure 4D compared to Figure 3A). In contrast, knocking down either *upd1* or *upd2* had no effect (Figure 4D). Use of cell specific drivers revealed that *upd3* was only required in the MpT principal cells but not in the stellate cells (Figure S2A). We confirmed the role of MpT Upd3 in gut homeostasis by assessing JAK-STAT pathway activity in the gut using the *10XSTAT-GFP* reporter. As expected, silencing *upd3* using the *uro-Gal4* driver substantially reduced GFP signal in the gut compared to the control line. This effect was again observed at 4 hpi but not at 16 hpi, indicating that MpT Upd3 contributes to the early phase of epithelium renewal (Figure 4E).

**Figure 4.**
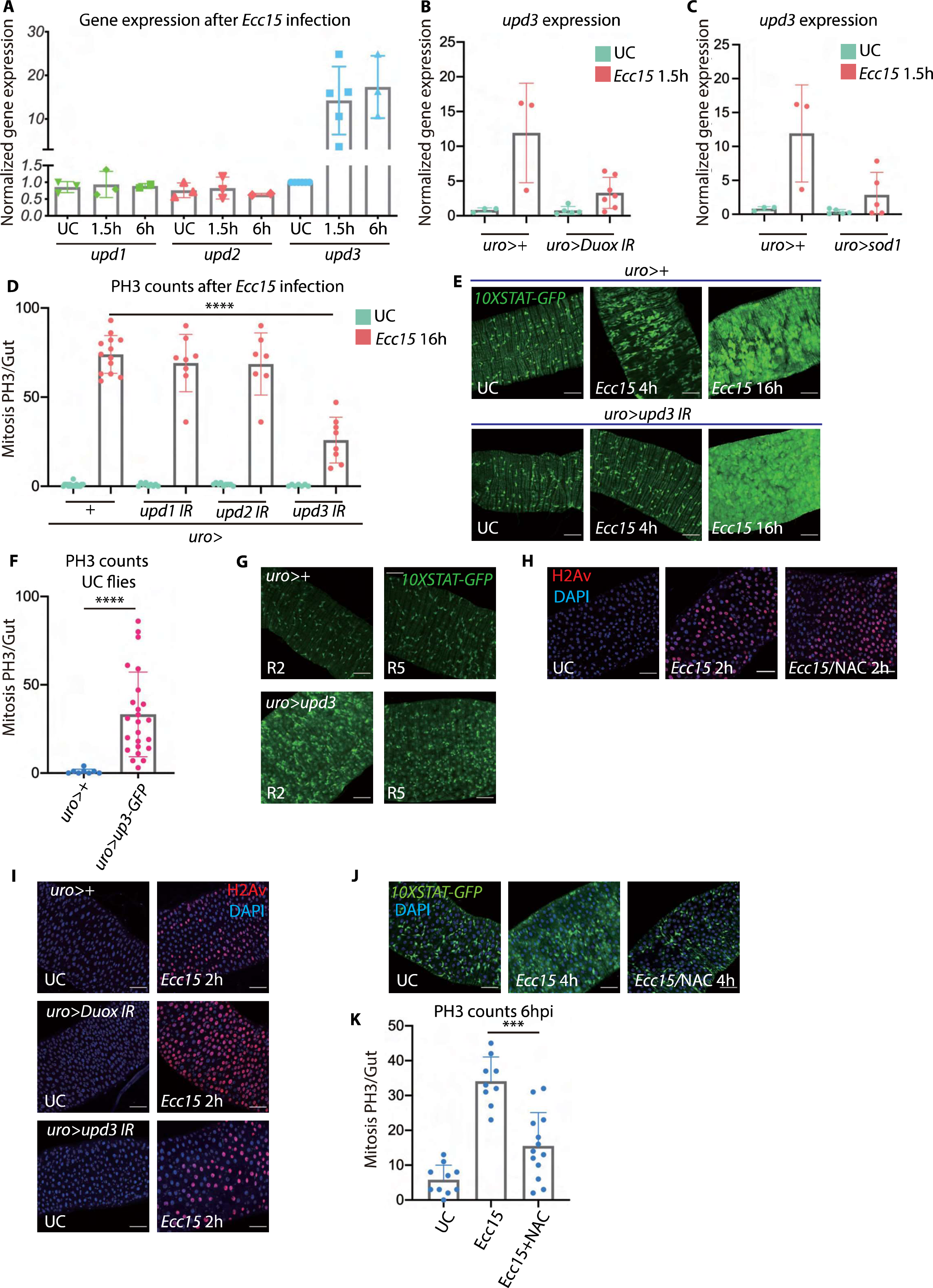
Tubules derived Upd3 but not ROS directly contributes to gut homeostasis after infection. **(A)** Only *upd3* is upregulated in MpT at early time points following *Ecc15* infection as revealed by RT-qPCR. (**B** and **C**) *upd3* expression in Malpighian tubules after oral infection required *Duox* (B) and is reduced in flies overexpressing the ROS scavenger *Sod1* (C). (**D**) Knockdown of *upd3* but not *upd1* or *upd2* in MpT impaired intestinal epithelial renewal as monitored by PH3 staining. (**E**) Silencing *upd3* in the MpT by RNAi reduced the increase JAK-STAT pathway observed 4 hpi but not at 16 hpi. (**F** and **G**) Over-expression of *upd3* in MpT increases the level of epithelial renewal in the gut as monitored by PH3 staining (F) and *10XSTAT-GFP* reporter activity (G). (**H**) The level of tissue damage in gut was monitored by H2Av staining in unchallenged and 2hpi *Ecc15* infected flies fed with 25mM of the antioxidant NAC. (**I**) Silencing *Duox* or *upd3* in the MpT did not impact the level of intestinal damage as shown by H2Av staining. (**J-K**) Feeding flies with NAC reduced the level of epithelial renewal after *Ecc15* oral infection as revealed by the *10XSTAT-GFP* reporter activity (J) and PH3 positive cell count (K). *** for *P* between 0.0001 and 0.001, **** for *P*≤0.0001 (F and K, Mann-Whitney test; D, one way ANOVA).

Overexpression of *upd3* in the principal cells (genotype *uro-Gal4>uas-upd3*) was sufficient to activate stem cell mitotic activity and increase *10XSTAT-GFP* expression level, confirming that Upd3 in MpT can remotely activate the stem cell renewal program in the midgut (Figure 4F and G). Surprisingly, the impact of Upd3 on intestinal stem cell proliferation was not restricted to the gut R5 region adjacent to the midgut-hindgut boundaries where MpT connect to the gut, but was observed all along the midgut including the R2 anterior region (Buchon et al., 2013b) (Figure 4G).

### Duox-produced ROS does not contribute to DNA damage upon infection

It is generally believed that damage caused by Duox-derived microbicidal ROS produced during gut infection cause stem cell compensatory proliferation (Buchon et al., 2009b, 2013a; Kim and Lee, 2014). However, we show instead that Duox is required in the principal cells of the MpT to produce ROS requiring for the production of Upd3, which stimulates epithelial renewal in the gut. We therefore asked whether ROS also directly contributes to gut epithelial renewal by causing damage, independent of Upd3 signaling. To monitor epithelial damage in the gut, we used H2Av staining to detect double-strand DNA breaks following *Ecc15* feeding. *Ecc15* feeding strongly induced DNA damage in the *Drosophila* gut at early time points (2hpi), particularly in the R2 anterior region of the midgut (Figure 4H). We then co-fed the flies *Ecc15* with N-Acetyl Cysteine (NAC), an antioxidant that decreases whole-body ROS upon feeding. However, decreasing ROS activity with NAC did not reduce DNA damage in the gut (Figure 4H), suggesting that ROS does not appreciably contribute to DNA damage at early time points during infection. Furthermore, knocking down *Duox* or *upd3* in MpT did not reduce the level of DNA damage at 2hpi with *Ecc15* (Figure 4I), indicating that Duox-produced ROS or Upd3-induced epithelial renewal do not contribute to DNA damage upon infection. However, feeding NAC did effectively reduce epithelial renewal as shown by decreased numbers of mitotic stem cells and JAK-STAT activity in the gut early in infection (Figure 4J and K). Interestingly, feeding flies with NAC reduced Upd3 expression in the MpT upon *Ecc15* gut infection (Figure S2B). This indicates that ROS do not primarily contribute to epithelial renewal by inducing damage but rather by impacting Upd3 expression in the MpT downstream of Duox. Collectively, these data show that Upd3 is the signaling molecule in the MpT downstream of Duox that stimulates the early phase of epithelial renewal of the intestine upon infection.

### A countercurrent flow flushes Upd3 along the gut

Early stimulation of epithelium renewal is not localized to the gut-MpT junction but spans the entire midgut including the anterior regions (Figure 4E). This suggests that MpT-derived Upd3 has a long-range action. Upd3 produced by MpT could reach the midgut epithelium either through the hemolymph (the insect blood) or via the gut-MpT connection (referred to as the luminal route). To distinguish between these two routes, we analyzed the fate of Upd3 produced by MpT. Western blot of flies over-expressing a *Upd3-GFP* fusion protein in MpT (*uro-GAL4>uas-upd3-GFP*) revealed the presence of Upd3 in the midgut but not in the hemolymph (Figure 5A). This observation strongly suggests that Upd3 reaches the gut through the luminal route. As MpT connect at the posterior end of the gut, the bolus flow generated by peristalsis is expected to push Upd3 towards the hindgut and not the midgut.

**Figure 5.**
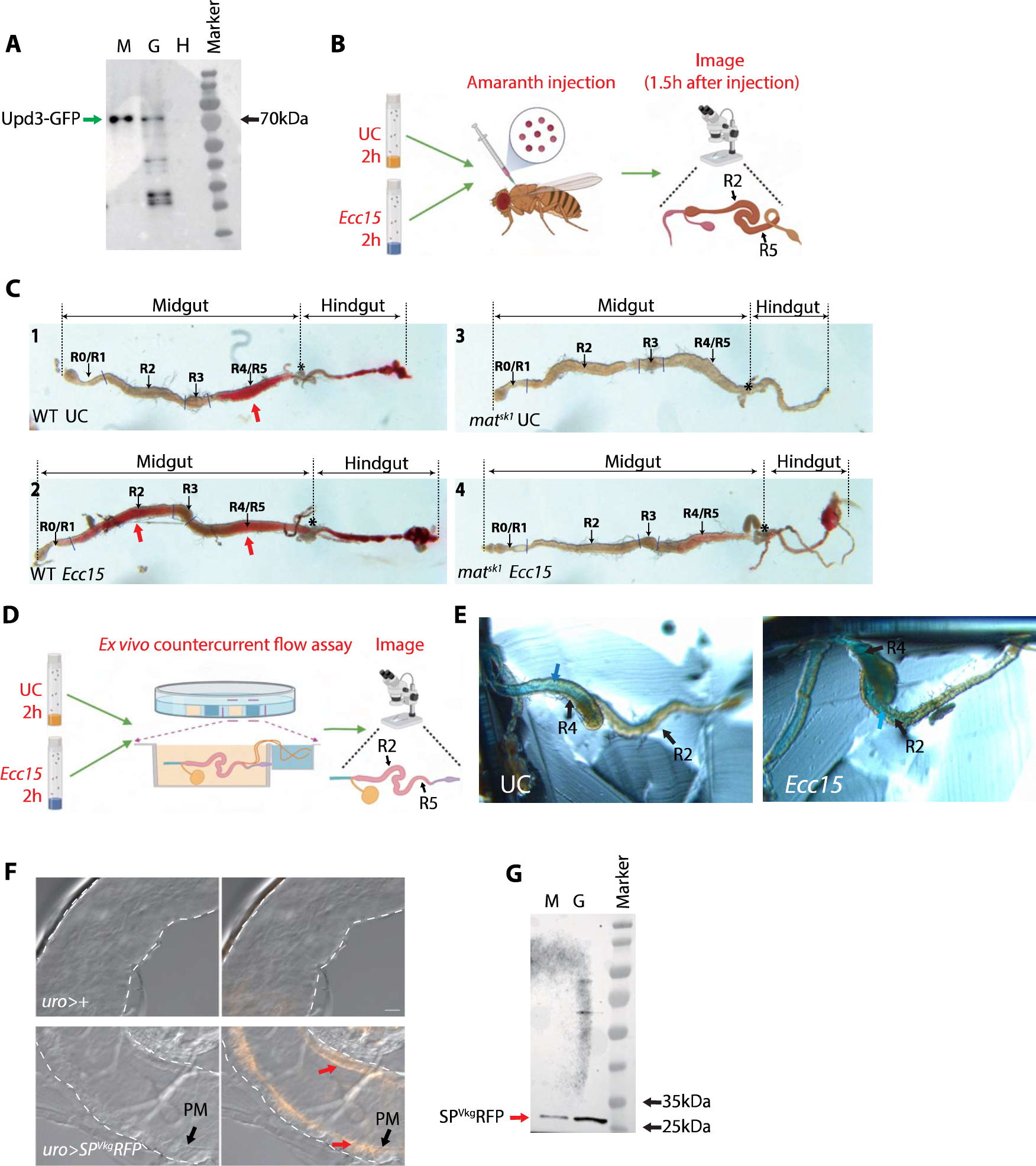
Malpighian tubules mediate countercurrent flow in the gut (**A**) Western blot monitoring the presence of *Upd3-GFP* in the MpT (lane M), midgut (lane G) and hemolymph (lane H), of flies over-expressing a *upd3-GFP* construct in the MpT (*uro- Gal4>uas-upd3-GFP)*. Green arrow indicates Upd3-GFP. (**B**) Schematic diagram of the experimental setup used for *in vivo* monitoring of countercurrent flow. UC: unchallenged. (**C**) Representative picture showing the presence of amaranth dye in the gut of wild-type and *mat^sk1^* flies either unchallenged or after *Ecc15* oral infection. In wild type unchallenged flies, the amaranth dye is observed in posterior midgut (R4 & R5 regions). A stronger amaranth accumulation is observed indicated by amaranth accumulation all along the midgut till R2 region in flies collected 2h after *Ecc15* oral infection. No strong amaranth accumulation was observed in the gut of *mat^sk1^* confirming that dye accumulation is not due to gut absorption. UC: unchallenged. * indicates the midgut-hindgut junction where MpT are connected. Red arrow indicates accumulation of Amaranth dye. (**D**) Schematic diagram of the experimental setup for *ex vivo* monitoring of the countercurrent flow. A two-well experimental PDMS plate setup was used to monitor countercurrent flow from MpT to the gut. UC: unchallenged. (**E**) Representative pictures showing the transfer of FD&C blue dye from the MpT to the gut lumen (blue arrow). (**F** and **G**) Expression of a secreted form of RFP in the MpT (*uro- Gal4>uas-SP^Vkg^RFP*) revealed the presence of the protein in the midgut lumen as revealed by live picture (F) and western blot (G) in flies collected 3h after *Ecc15* oral infection. Red arrows indicate the presence of RFP. PM: Peritrophic matrix.

The MpT-gut luminal route would therefore require a countercurrent flow to flush Upd3 backwards, from the midgut-hindgut junction to the anterior midgut. The existence of such countercurrent flow has been described in the gut of several insect species, notably locusts, but not in Diptera species like *Drosophila* (Dow, 1981; Terra and Ferreira, 2012). To further elucidate the mechanism underlying MpT-gut communication, we explored the hypothesis of a countercurrent flow system in *Drosophila*. To this end, we used amaranth, a dye which is efficiently absorbed by MpT but minimally absorbed by the gut (Dow, 1981). We injected amaranth into the body cavity and allowed absorption by MpT (Figure 5B). We then dissected the fly gut to check whether the dye was pushed towards the anterior part of the midgut. We observed the presence of the dye in the posterior region R4 and R5 of midgut of unchallenged flies (Figures 5C1). Interestingly, we found that oral bacterial infection induced a much stronger countercurrent flow compared to the unchallenged condition, as shown by stronger accumulation of dye along the midgut upward to the R2 region of the anterior midgut, where a high level of epithelium renewal is observed upon infection (compare Figure 5C2 to 5C1). We did not observe the strong presence of the dye in the upper anterior midgut region R0 and R1. To confirm that the accumulation of dye in the gut was indeed caused by a countercurrent flow originating from MpT, we repeated this experiment in *mat^sk1^* mutant flies. Materazzi (Mat) is a lipid binding protein that protects MpT from ROS during stress and infection (Li et al., 2020). We have previously shown that *mat* deficient flies accumulate ROS in the MpT, leading to tubule dysfunction and reduced excretion (Li et al., 2020). Dye did not accumulate in the gut of *mat^sk1^* flies using our amaranth injection method, demonstrating that presence of dye in the midgut is not due to absorption through the intestine but rather by countercurrent flow originating from MpT (Figure 5C3 and C4). To provide a more direct demonstration of this countercurrent flow, we developed an *ex vivo* assay inspired by the Ramsay assay commonly used to monitor MpT secretion ability (Dow et al., 1994a). In this experiment, we placed the extremity of MpT in wells containing Schneider’s Insect Medium, insect saline buffer with FD&C blue dye (the dye well), while the gut was placed in another well filled with medium only. This allowed direct observation of dye flowing from the MpT to the midgut (Figure 5D). This *ex vivo* assay reveals the passage of blue dye from the MpT to the midgut, the effect being more marked when orally infected flies were used (Figure 5E). To further confirm the existence of this countercurrent flow, we overexpressed a secreted version of RFP in the MpT and monitored the presence of the protein in the gut lumen. Both live RFP detection and western blot confirm the presence of MpT derived RFP in the gut, consistent with the countercurrent flow hypothesis (Figure 5F and 5G). Interestingly, the secreted RFP protein was found in the ectoperitrophic space between the peritrophic matrix and the epithelial cells (Figure 5F).

### Countercurrent flow and epithelial renewal in the gut requires the aquaporin Drip

We then investigated the mechanism by which MpT generate the countercurrent flow. We hypothesized that this countercurrent flow would require water uptake by the tubules. Water influx in MpT is promoted by two aquaporins, the apical membrane protein Drip and the basal membrane protein Prip, both expressed in stellate cells (Cabrero et al., 2020). To test whether these water channels are involved in the countercurrent flow, we knocked down *Drip* and *Prip* by RNAi using the stellate cell *Gal4* driver *c724-Gal4*, and monitored countercurrent flow by injection of amaranth as described in Figure 5C. Flies with reduced

*Drip* expression exhibited a marked reduction in countercurrent flow, particularly upon *Ecc15* infection (Figure 6A). In contrast silencing *Prip* did not reduce countercurrent flow as indicated by the presence of amaranth signal in the gut. These data indicate that the aquaporin Drip contribute to countercurrent flow activity, likely by promoting transfer of water from the hemolymph to the MpT lumen.

**Figure 6.**
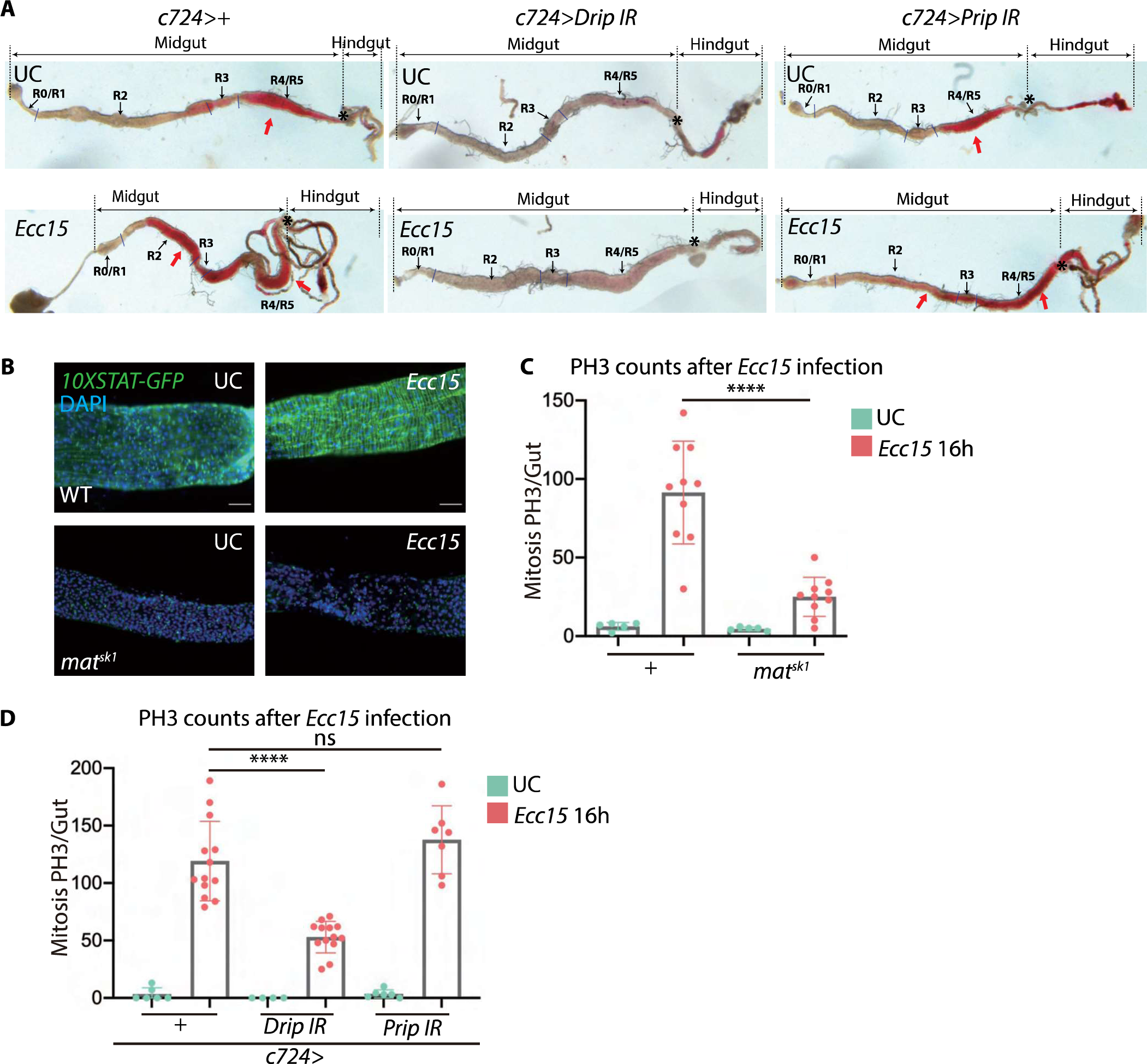
Regulation of countercurrent flow by an aquaporin (**A**) The countercurrent flow was monitored using the amaranth injection assay in wild-type flies and aquaporin knockdown flies (*c724-Gal4>uas-Prip IR* and *c724-Gal4>uas-Drip IR*) either unchallenged or collected 2hpi. UC: unchallenged; * indicates midgut-hindgut junction where the MpT are connected. Red arrow indicates accumulation of Amaranth dye. (**B** and **C**) The level of intestinal epithelial renewal is reduced in *mat^sk1^* flies as shown by reduced JAK-STAT pathway activity (B) and low PH3 postive cell count (**C**). (**D**) The level of epithelial renewal is reduced when *Drip* is silenced in stellate cells. UC: unchallenged flies. **** for *P*≤0.0001, ns: non-significant (C, Mann-Whitney test; D, one way ANOVA).

The existence of a countercurrent flow likely explains the ability of MpT to remotely contribute to intestinal homeostasis. To directly test this notion, we monitored the level of epithelium renewal upon oral bacterial infection in *mat* and *Drip* deficient flies with defective countercurrent flow. Consistent with our model, we observed that *mat^sk1^* flies have reduced intestinal epithelium turnover and reduced JAK-STAT activity in the gut (Figure 6B and C). This indicates that functional MpT are critical for countercurrent flow generation and intestinal homeostasis. Similarly, silencing *Drip* in the stellate cells led to a strong decrease in the level of stem cell proliferation after *Ecc15* oral infection (Figure 6D). We conclude that the countercurrent flow generated by MpT excretory function contributes to intestinal homeostasis.

## Discussion

### Duox is required in the MpT to simulate intestinal epithelial renewal

Although the role of Duox in intestinal immunity has been well established, our study challenged the common view by showing that this NADPH oxidase is required in the MpT, and not in the gut as originally thought. Previously, Ha et al. demonstrated that Duox is required in the gut for generating microbicidal ROS that contribute to the elimination of ingested bacteria. In their study, they used the ubiquitous driver *da-Gal4* and the posterior midgut driver *cad-Gal4* to silence *Duox* (Ha et al., 2005a). However, high-throughput sequencing data clearly show that *cad* is highly enriched in MpT (Leader et al., 2018).

Furthermore, *cad>GFP* flies not only express GFP in the posterior midgut, but even more strongly in the lower ureter of MpT (Buchon et al., 2013b; Ryu et al., 2008)(L.X. unpublished observation). Buchon et al. later revealed that knockdown of Duox impaired gut epithelial renewal, raising the hypothesis that epithelium renewal was a consequence of ROS-mediated damage (Buchon et al., 2009b). However, their observations were also based on the use of ubiquitous driver *da-Gal4* to knockdown *Duox*. Here we used a combination of gut and MpT specific drivers to reveal that the impact of Duox on gut homeostasis is due to Duox activity in MpT. Consistent with this, Duox is expressed at a much higher level in MpT compared to the gut, and Duox-derived ROS products are generated in higher amounts in MpT. Use of three MpT cell-specific drivers (*uro-Gal4*, *capaR-Gal4* and *c42-Gal4*) and another *Duox* RNAi construct confirmed that Duox is required in the principal cells of the tubules to stimulate intestinal stem cell proliferation. Collectively, our study confirms that Duox contributes to the early phase of epithelium renewal, but probably not as a consequence of tissue damage through ROS. Consistent with this, we did not observe a ROS burst in the gut at the early time point after infection. Jones et al. also found that *Ecc15* ingestion did not increase ROS in the *Drosophila* gut at 2hpi (Jones et al., 2013). Moreover, we found that reducing ROS with antioxidant did not reduce the level of DNA damage in the midgut upon *Ecc15* infection. Thus, ROS may not be the direct cause of tissue damage upon infection.

This is in line with the observation that enterocyte delamination induced upon gut infection does not require Duox (Zhai et al., 2018). Besides inducing a ROS burst, pathogenic bacteria release a wide range of toxins that could directly damage epithelial cells in both Drosophila and mammals (Mileto et al., 2020; Nehme et al., 2007; O’Brien and Holmes, 1987; Opota et al., 2011; Vallet-Gely et al., 2008).

The role of ROS in general and NADPH oxidases in particular in gut immunity remains poorly characterized. Studies in *Drosophila* and insects point to a role of ROS as a signaling molecule (this study and Jones et al., 2013), a microbicidal agent (Ha et al., 2005a), and sclerotization agent (Jang et al., 2021; Kumar et al., 2010). In mammals, Nox1 and Duox 2 are strongly expressed in intestinal epithelial cells (Lambeth and Neish, 2014). ROS produced by NADPH oxidases serves an antimicrobial function and is involved in the maintenance of microbiota homeostasis (Corcionivoschi et al., 2012; Geiszt et al., 2003; Kumar et al., 2007; Matziouridou et al., 2018). However, other NADPH oxidases, notably Nox1, Nox4 and Nox5, also exert a signaling function (Lambeth and Neish, 2014). Thus, NADPH oxidases-mediated ROS in the intestine appears to have multiple roles in gut homeostasis that are far from being well understood. By showing that Duox in the MpT regulates intestinal homeostasis, we revealed that this NADPH oxidase can have long-distance effect, in line with other studies involving Duox in hemocytes or trachea (Chakrabarti and Visweswariah, 2020; Jang et al., 2021).

### Duox functions upstream of Upd3 to simulate intestinal epithelial renewal

A recent study showed that ROS produced by Duox in hemocytes leads to the expression of the cytokine Upd3 (Chakrabarti and Visweswariah, 2020). Similarly, we observed that Duox activity regulates the production of Upd3 in MpT to promote the early phase of epithelium renewal. Thus, both studies highlight the role of Duox-mediated ROS as a signaling molecule upstream of Upd3. Previous studies did not identify the ROS species required for *upd3* induction (Gordon et al., 2018; Santabárbara-Ruiz et al., 2015; Srinivasan et al., 2016). Our observation that overexpression of superoxide dismutase Sod1 but not the extracellular catalase Irc or Cat blocks *upd3* induction suggests that superoxide is the ROS species responsible for *upd3* induction. The pathways leading to activation of Duox in the MpT by bacterial oral infection and linking ROS to Upd3 expression in the MpT are currently unclear. Other studies have suggested that *upd3* induction by ROS requires the p38 and JNK pathways in imaginal discs and the Src42A/Shark/Draper pathway in hemocytes (Chakrabarti and Visweswariah, 2020; Santabárbara-Ruiz et al., 2015). In the midgut, *upd3* has been shown to be regulated by the Hippo, TGF-β, and Src-MAPK pathways in the context of immune challenge (Houtz et al., 2017). Future studies should further characterize the regulation of *upd3* by Duox in MpT and investigate a possible role for the components described above. Upds are also produced by intestinal cells, notably the stem cells, enteroblasts and enterocytes, to regulate the level of epithelial renewal (Jiang et al., 2009). We therefore wonder why Upd3 production is required in the MpT to promote the early phase of intestinal epithelial renewal. An attractive hypothesis is that rapid release of Upd3 by MpT could boost the early phase of midgut epithelial renewal when intestinal cells are under assault by damaging agents.

### Evidence for a countercurrent flow in *Drosophila*

The existence of a countercurrent flow in the digestive tract of insects has been described in several species. In 1933, Wigglesworth had already observed that Trypan blue fed to mosquitoes accumulates in the anterior part of the gut. This led him to propose that fluid in the gut lumen must circulate from the posterior to the anterior part of the gut (Wigglesworth, 1933). In 1970, Berridge established a countercurrent flow model, showing that fluid secreted from the posterior to the anterior midgut created an absorption cycle with a role in food digestion (Berridge, 1970). Dow observed that countercurrent flow was generated by MpT of starved but not fed locusts, showing that this current can be regulated (Dow, 1981). The existence of retrograde fluid flow in the insect gut is possible due to the presence of a peritrophic matrix that transversally compartmentalizes the midgut (Terra and Ferreira, 2012). To our knowledge, countercurrent flow has never been demonstrated in *Drosophila* despite ever-growing interest in gut physiology. By monitoring injected dyes and secreted proteins, we show that a countercurrent flow also exists in this insect from midgut-hindgut adjacent region where MpT connect to the gut, to the anterior part of the midgut. The observation that *mat^sk1^* flies with defective MpT function did not exhibit countercurrent flow confirms that it originates from MpT. Stellate cells of MpT provide a route for rapid water efflux into *Drosophila* renal tubules through the aquaporins Drip and Prip (Cabrero et al., 2020). Our study showed that the countercurrent flow relies on the water channel Drip in stellate cells. In insects, MpT activity is largely controlled by diuretic hormones (Chapman et al., 2013). Tyramine and leucokinin are two main diuretic hormones regulating stellate cell activity (Cabrero et al., 2013). Future studies should investigate the role of neuropeptides in the regulation of this countercurrent flow. Use of amaranth dye reveals that the *Drosophila* countercurrent flow only reached the R2 region, where the staining ends abruptly. This suggests an active absorption process in this region. Interestingly, the R2 region is highly enriched with aquaporins, notably Drip (Buchon et al., 2013b), that could contribute to water reabsorption. It is fascinating to note that one of the MpT pairs directed upward are attached to the midgut at this R2 region. Although this positioning may simply extend the MpT upward to favor the cleaning of the whole-body cavity, an enticing hypothesis is that this orientation allows direct transfer of water absorbed in the R2 midgut region to the MpT, establishing a MpT-gut-MpT cycle. It should be noted that the R1 anterior region does not benefit from the protection provided by the counterflow. As Duox is also highly expressed in the crop and cardia, the R1 region may be protected by a flow coming from the foregut, a hypothesis that remains to be investigated.

### Critical role of MpT in intestinal immunity

Although almost 90 years have passed since its discovery, the function and molecular mechanism of the countercurrent flow system is still poorly understood. The most well-established role of countercurrent flow is in promoting enzyme recycling (Terra and Ferreira, 1981). As the food bolus moves toward the posterior midgut, nutrients and digestive enzymes pass through the peritrophic matrix into ectoperitrophic space where countercurrent flow flushes them toward the anterior midgut (Hegedus et al., 2009). In this study, we found that oral infection strongly increases countercurrent flow activity in the gut, pointing to its role as a host defense mechanism. We also show that this countercurrent flow sustains the early phase of epithelial renewal through Upd3 to ensure barrier integrity. The contribution of MpT to host defense is likely not restricted to epithelial renewal. MpT are potent immune reactive organs producing antimicrobial peptides (AMPs) and other immune molecules in response to infection (Kaneko et al., 2006; Tzou et al., 2000; Zugasti et al., 2020). Thus, MpT may also facilitate microbial clearance by flushing antimicrobials into the gut via countercurrent flow. The countercurrent flow provides a fascinating mechanism to delegate production of antimicrobial factors to MpT. This would allow the transfer of antimicrobials to the gut even if gut function has been impaired. This process could be important in preventing intestinal infections that are likely to be frequent in insects such as *Drosophila* that feed on rotting fruits. We were impressed by the strength and rapidity of the counterflow current. MpT is considered the most potent epithelium for fluid production on a per cell basis (Beyenbach et al., 2010; Maddrell, 1991). Thus, MpT provide adequate fluid transfer to allow a rapid circulation within the midgut, possibly removing toxins and introducing microbials.

Collectively our study highlights the critical role of MpT in gut immunity, revealing a striking example of inter-organ communication. There is no evidence for countercurrent flow in the mammalian intestine so far. The existence of localized fluid flow cannot however be excluded, as fluids are secreted at the level of the crypt and absorbed at the tip of villi (Kiela and Ghishan, 2016). Thus, local flow could establish a gradient of antimicrobial molecules from the crypt to the lumen. This shows that regional or local fluid fluxes are likely an important aspect of intestinal homeostasis and immunity in all animals, which requires further investigation.

## Acknowledgement

We thank the Bloomington Stock center, the Vienna *Drosophila* Resource Center and National Institute of Genetics (NIG) for fly stocks. We thank Hannah Westlake, Florent Masson and Samuel Rommelaere for careful reading and comments on the manuscript. We thank Emőke Gerőcz for assistance in some experiments. We would also like to thank Jean-Philippe Parvy, Won Jae Lee and Jun Zhou for kindly providing fly stocks. This work was supported in part using the resources and services of the BioImaging and Optics Platform Research Core Facility at the School of Life Sciences of EPFL. The schematic diagrams are created with Biorender.com.

## Author contributions

X.L. and B.L. designed the study. X.L. conducted the experiments. X.L. and B.L. interpreted the data and wrote the manuscript.

## Supplementary Figure Legends

**Figure S1. Duox is required in MpT to simulate intestinal epithelial renewal upon infection (related to** Figure 3) (**A**) The level of epithelial renewal as monitored by PH3-postive cells count is reduced 14hpi following *Ecc15* infection when *Duox* is knockdown in MpT (*capa-Gal4>Duox IR*). (**B**) The level of epithelial renewal as monitored by PH3-positive cells count is reduced 16hpi following *Ecc15* infection when *Duox* is knockdown in MpT confirmed by another Duox IR construct (*uro-Gal4>Duox IR* (BL38916)). (**C**) Tissue specific *Duox* knockout by CRISPR-Cas9 in MpT principal cells decrease epithelial renewal as revealed by PH3 staining. ** for *P* between 0.001 and 0.01, (A and C, Mann-Whitney test; B, one way ANOVA).

**Figure S2. Tubules derived Upd3 but not ROS directly contributes to gut homeostasis after infection (related to** **Figure 4**). (**A**) Silencing *upd3* in MpT stellate cell (*c724-Gal4>upd3 IR*) do not alter the level of epithelial renewal as monitored by PH3-positive cells count after *Ecc15* infection. (**B**) Feeding flies antioxidant NAC reduces *upd3* expression in MpT. ns: non-significant, Mann-Whitney test.

