## Supplementary figures and images for "Duox activation in *Drosophila* Malpighian tubules stimulates intestinal epithelial renewal through a countercurrent flow"

### Supplemental Figures 1 and 2

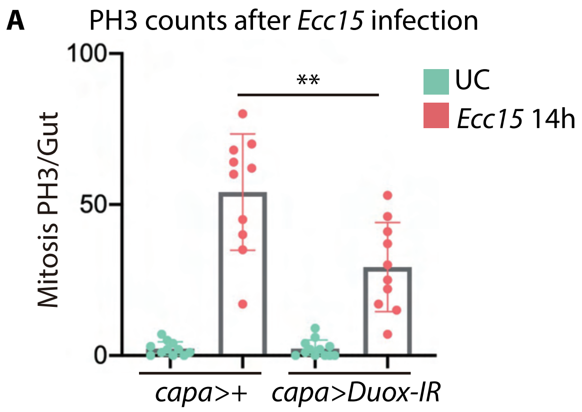

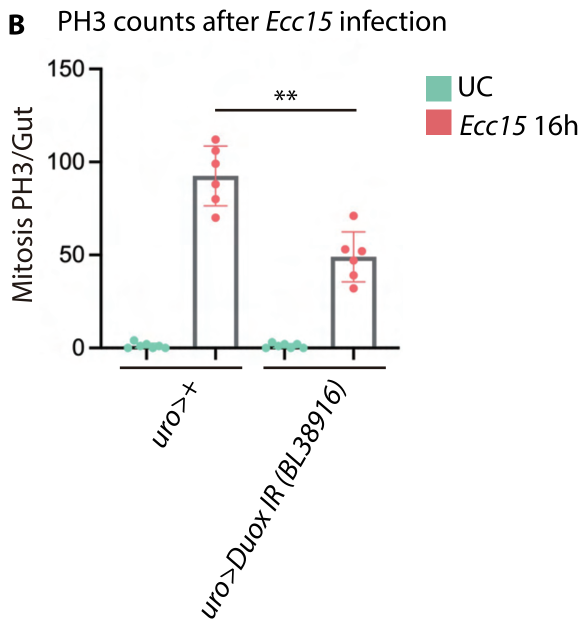

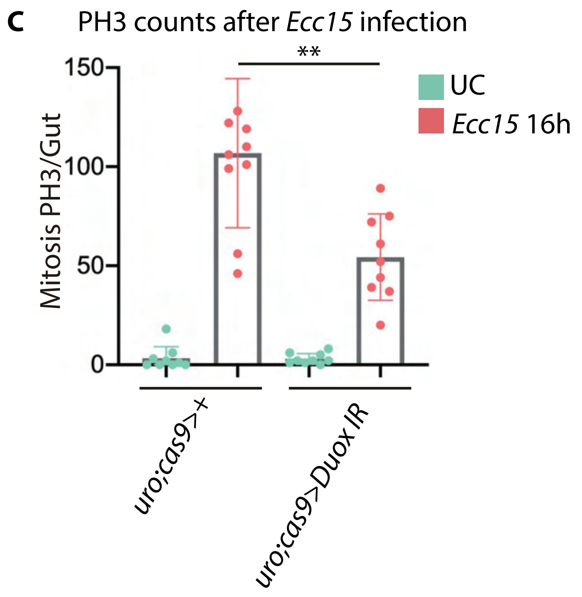

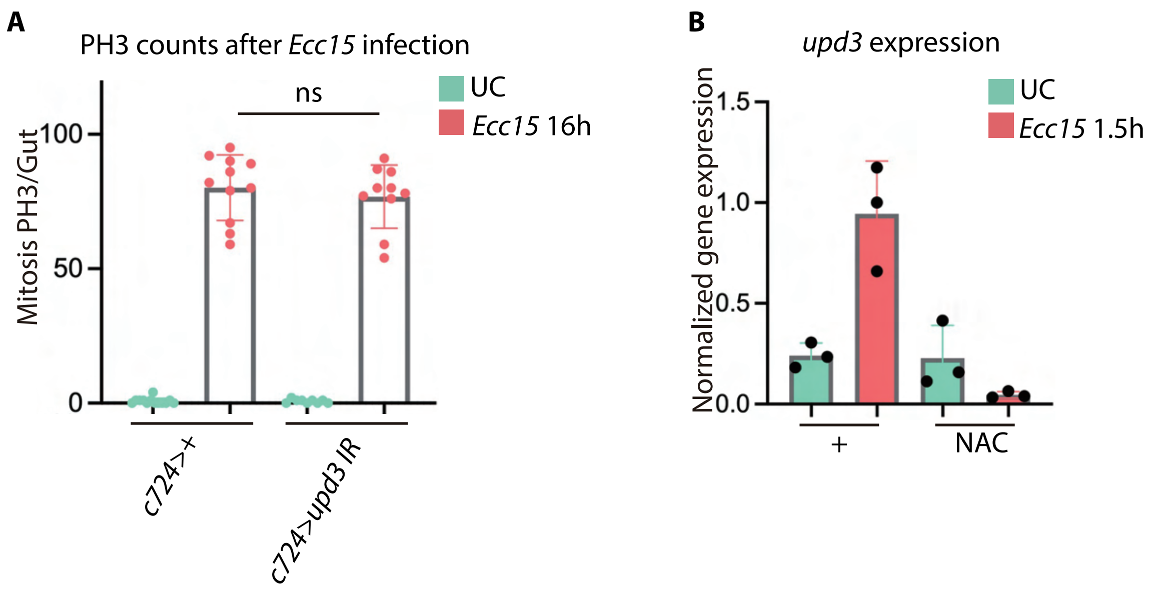
